# Disruption of IL-17 signaling in the respiratory mucosa results in invasive streptococcal infection

**DOI:** 10.1101/2023.11.10.566548

**Authors:** Jamie-Lee Mills, Ailin Lepletier, Victoria Ozberk, Jessica Dooley, Jacqualine Kaden, Ainslie Calcutt, Yongbao Huo, Allan Hicks, Ali Zaid, Michael F. Good, Manisha Pandey

## Abstract

*Streptococcus pyogenes* infection of the upper respiratory tract and skin can lead to severe invasive streptococcal disease (ISD). Previous studies have demonstrated that the deficiency of IL-17 in mice (IL-17-/-) reduces mucosal immunity against *S. pyogenes*. However, the impact of IL-17 deficiency on the development of ISD is unknown. Here, we model single or repeated non-lethal, intranasal (IN) *S. pyogenes* M1 strain infections in immunocompetent and IL-17-/-mice to assess bacterial dissemination following a final IN or skin challenge. Immunocompetent mice that received a single *S. pyogenes* IN infection displayed long-lasting mucosal immunity and no systemic infection. However, in the absence of IL-17, a single IN infection resulted in the dissemination of *S. pyogenes* to the spleens, which was further exacerbated by repeated IN infections. Interestingly, immunity following skin challenge did not show a correlation with IL-17 and was instead associated with the activation of germinal center responses and the accumulation of neutrophils in the spleen. Our results highlight the critical role of IL-17 in preventing ISD following *S. pyogenes* infection of the respiratory mucosa.

## INTRODUCTION

*Streptococcus pyogenes* (*S. pyogenes*) colonizes the upper respiratory tract and skin. In some cases, mucosal and skin barriers become vulnerable to bacterial escape, leading to invasive infections. These infections include potentially life-threatening conditions such as sepsis, pneumonia, necrotising fasciitis and toxic shock syndrome. *S. pyogenes* infection can also lead to post-streptococcal autoimmune diseases, primarily acute rheumatic fever and rheumatic heart disease. Collectively, streptococcus-related pathology is responsible for the loss of approximately 500,000 lives each year (Carapetis, Steer et al., 2005, Nelson, Pondo et al., 2016), with the greatest burden suffered by people in developing countries and Indigenous populations living in economically advanced societies (Bocking, Matsumoto et al., 2016). Natural immunity to *S. pyogenes* at the primary site of infection is slow to develop and a vaccine is not available.

Antibodies (IgA and IgG) (Mortensen, Christensen et al., 2017, Mortensen, Nissen et al., 2015, Pandey, Ozberk et al., 2016a), effector immune cells (CD4^+^ T cells, macrophages and neutrophils) (Dileepan, Linehan et al., 2011, Hyland, Brennan et al., 2009, Mishalian, Ordan et al., 2011) and cytokines (including IL-17A and IFN-ɣ) (Dileepan et al., 2011, Edwards, Taylor et al., 2005, Hyland et al., 2009) have been shown to play critical roles in regulating immune responses to *S. pyogenes* at the site of infection. These immune responses can collectively target multiple streptococcal antigens, with a key antigen being the major virulence factor, the M-protein (encoded by the *emm* gene). In addition to the M-protein, other bacterial virulence factors have also been shown to suppress innate and acquired immune responses to *S. pyogenes* (Walker, Barnett et al., 2014). However, the immune mechanisms that facilitate systemic dissemination of *S. pyogenes* from the respiratory mucosa and skin are elusive.

Early streptococcal research in humans demonstrated that protection against homologous strains following pharyngeal infection is long-lasting. M-type-specific antibodies were recovered in convalescent blood following pharyngitis (Kuttner & Lenert, 1944, Rothbard, 1945), with bacteriostatic properties persisting in some individuals (Rothbard, 1945) and remaining present in the blood for a substantial length of time, with the longest duration reported by Lancefield being up to 32 years (Lancefield, 1959). While earlier studies focused on infections of the upper respiratory tract (URT), later studies in First-Nation Australian communities, where skin infection is far more prevalent than pharyngitis, identified different strains moving through communities. In some cases, more than one strain existed at a time and persisted for longer than 6 months (McDonald, Towers et al., 2007b). Thus, it is likely that *S. pyogenes* strains will remain in a community due to a reservoir of skin infection. In tropical communities with high rates of rheumatic heart disease, immunity to a skin strain of *S. pyogenes* was clearly slow to develop (McDonald et al., 2007b). In agreement with these clinical findings, our previous study using a murine model of invasive disease associated with skin pyoderma, show that immunity at the skin and systemic sites required repeated homologous skin infections (Pandey et al., 2016a). In that study, enduring protection correlated with M-type-specific memory B cell responses in the sera, spleen and bone marrow, in the absence of which immunity was rapidly lost (Pandey et al., 2016a).

While systemic immunity to *S. pyogenes* infection is associated with IgG responses in the sera, mucosal immunity relies on the production of secretory IgA (which can prevent the attachment of streptococcus to mucosal surfaces (Fluckiger, Jones et al., 1998)), and IL-17 (which results in recruitment of neutrophils and other inflammatory cells that contribute to bacterial clearance at mucosal sites (Carey, Weinberg et al., 2016, Lu, Gross et al., 2008). Accordingly, inborn IL-17 deficiency and therapeutics based on IL-17 inhibitors increase the risk of mucocutaneous candidiasis in humans (Davidson, van den Reek et al., 2022, Puel, Cypowyj et al., 2011, Puel, Cypowyj et al., 2012). Similarly, the most common adverse effects of anti-IL-17 therapy in patients with psoriasis is an increase in infections of the nasopharyngeal tract (Wang, Wang et al., 2023).

Besides *S. pyogenes*, the route of infection by other bacteria can cause a fundamental difference in the resulting immune responses (Hu & Pasare, 2013). It has been observed that intranasal *Francisella tularensis* infection induces a Th17 response in the lungs, whereas the intradermal route of infection favours a Th1 response in both spleens and lungs (Woolard, Hensley et al., 2008). Therefore, immunity in the mucosal, skin and systemic sites may be regulated separately. These diverse immune responses may impede the development of resistance to infection in those cases where the organism can infect via different anatomical sites.

In this study, we investigate the development of mucosal and systemic immunity following single or multiple URT infections with a homologous *S. pyogenes* isolate and correlate it with immune mechanisms underpinning site-specific or cross-compartmental protection. Through the assessment of both humoral and cellular immune responses and via the use of IL-17 deficient mice, we show that CD4^+^ T cells, IL-17 and neutrophil responses in the lungs collectively regulate protection at the respiratory mucosa. This is the first study to show that in the absence of IL-17, URT infections result in the passage of *S. pyogenes* from mucosal sites into lymph nodes and spleen. This research provides critical insight into the role of IL-17 in orchestrating the interplay between immune cells in the respiratory mucosa and systemic sites, highlighting the importance of integrating strategies that are capable of inducing IL-17 responses, alongside humoral responses, in the design of vaccines.

## RESULTS

### A single *S. pyogenes* IN infection results in enduring immunity at the respiratory tract

We initially sought to demonstrate immunity in the respiratory tract following intranasal (IN) infection. Cohorts of BALB/c mice received either 1 or 2 non-lethal IN infections with mouse-passaged *S. pyogenes* 2031 (*emm*1), each 3 weeks apart (Fig 1A). To assess if the number of prior intranasal exposures would determine the level of site-specific protection, mice received a final IN challenge with homologous *S. pyogenes* isolate three weeks following a single or two sequential infections. Mice were closely monitored for the appearance of clinical symptoms and scored based on an approved clinical scoring system (Ozberk, Reynolds et al., 2021b). All mice, regardless of number of prior IN exposures (0x, 1x or 2x), showed a significant reduction (p<0.05-0.001) in clinical scores when compared to the control group, which was not infected prior to challenge (Fig 1 B). On day 2 post-challenge, mice were euthanized to assess bacterial burden (colony forming units [CFU]) in the nasal associated lymphoid tissue (NALT), and in the lungs, as an indicator of immunity in the upper respiratory tract (URT) and lower respiratory tract (LRT). Regardless of number of prior exposures, all mice had significant reductions in bacterial burden in NALT (66.6-92.6%; p<0.05) and lungs (90-100%; p<0.01) when compared to the control group (0x) (Fig 1 C and D, respectively). IN challenge of these mice resulted in very low to undetectable bacterial burden at systemic sites (including spleen and lymph nodes), which was comparable across all groups, with or without prior exposure (Fig 1E).

**Figure 1.**
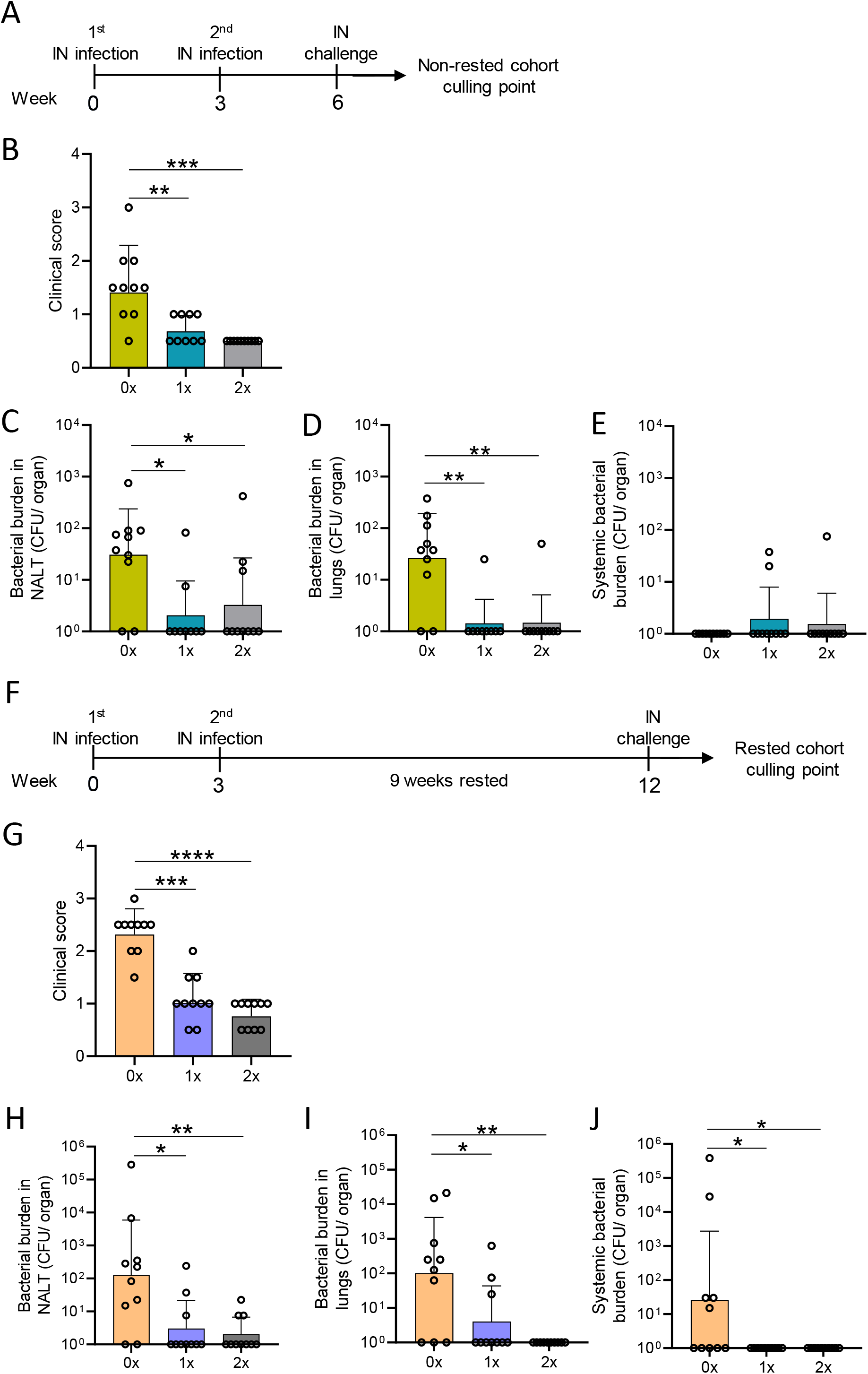
Assessment of mucosal protection and its endurance following 0-2 intranasal homologous infections, prior to IN challenge. (A) BALB/c mice (n= 10, female, 4-6 weeks old) were given IN infections with 2031 (*emm*1), each 3-weeks apart. Three weeks later, all mice received a homologous IN challenge. (B) on day 2 following challenge mice were assessed for clinical scores as per the approved score sheet. Mice were sacrificed on day 2 post IN challenge to assess bacterial load in (C) NALT, (D) lungs and (E) systemic sites (spleen and pooled cervical lymph nodes). (F) Alternatively, BALB/c mice (n= 10, female, 4-6 weeks old) given IN infections with 2031 were rested for 9 weeks before receiving a homologous IN challenge. (G) Clinical scores were assessed on day 1 and 2 following challenge and are shown as average. Mice were sacrificed on day 2 post IN challenge to assess bacterial load in (H) NALT, (I) lung, and (J) systemic sites (spleen and pooled cervical lymph nodes). Data shown are the CFU geometric mean ± geometric SD. Significance was determined using Mann-Whitney rank analysis of CFU in sequentially infected group compared to CFU of the challenge control (0x), * p<0.05, ** p<0.01, *** p<0.001, **** p<0.001. X-axis shows the number of infections prior to challenge. 1x only received infection at week 3 and 2x at both week 0 and 3. Mice were rested for 9 weeks and received challenged at week 12. 0x received challenge only.

To assess the longevity of protection at the respiratory tract, a separate cohort of mice (also previously infected 1 or 2 times with *S. pyogenes*) was rested for 9 weeks before receiving a homologous IN challenge (Fig 1F). We observed that single or two sequential infections (Fig 1 A-D) induced enduring immunity which did not wane following a 9 week rest. All mice previously infected with *S. pyogenes* demonstrated a significant reduction in clinical scores in comparison to the control mice (p<0.001-0.0001) (Fig 1G). This was associated with significantly reduced bacterial burden in NALT (96.3-99.4%, p<0.05-0.01) and lungs (93.8-99%, p<0.05-0.01) (Fig 1H and I, respectively). However, 2 IN infections were better for enduring immunity in the NALT and lungs. Interestingly, the IN challenge to mice not previously infected led to systemic infection only in the rested naïve mice (Fig 1J), which presented higher systemic burden than non-rested naïve mice (>99.9%, p<0.05), suggesting that age was associated with reduced innate protection. All mice with prior *S. pyogenes* infection demonstrated complete protection at the systemic sites (lymph node and spleen), which was significant in comparison to the control cohort (Fig 1J).

Overall, we found that a single IN exposure to *S. pyogenes* led to mucosal immunity at both the URT and LRT. Further, our data show an age-related susceptibility to bacterial systemic dissemination following *S. pyogenes* IN challenge; however this was completely prevented by previous exposure to the organism.

### Mucosal immunity against *S. pyogenes* at the respiratory tract is associated with local humoral and cellular responses

To investigate the role of humoral immunity in this mucosal protection, we measured the levels of M-protein type specific IgG in the serum of mice following each sequential infection. *S. pyogenes* M1 type-specific IgG antibody was not evident following a single IN infection (Supp Fig 1A). Nevertheless, M1-specific IgG titres developed after 2 infections and remained consistent in the cohorts that received subsequent IN infections (Supp Fig 1A).

The fact that M1-specific circulating antibodies were not evident in mice after a single *S. pyogenes* infection prompted us to investigate other immune correlates of protection at the respiratory mucosa. Salivary antibodies play important roles in protection from *S. pyogenes* by preventing bacterial attachment to the mucosal epithelia, opsonising the bacteria, and directly lysing them(Lamm, 1997, Markus Lilja, 1999). To investigate the role of URT mucosal immunity, we measured the levels of M1-specific IgG and IgA in the saliva of mice following each infection. M1-specific IgG and IgA titres were significantly increased in the saliva after a single infection (Fig 2A i & ii, respectively). A progressive increase in M1-specific IgG and IgA responses in the saliva with increased number of infections was also noted. To investigate long-term humoral immunity mediated by antibody secreting cells in the bone marrow (BM) (known as long-lived plasma cells (LLPCs)) (Brynjolfsson, Persson Berg et al., 2018, Nguyen, Joyner et al., 2019), we enumerated the number of M1-specific IgG-secreting LLPC in infected mice using an ELISpot assay. We found that the numbers of M1-specific IgG-and IgA-secreting BM cells were significantly higher in mice that received 2 IN infections than in naïve mice (Fig 2B i and ii, respectively).

**Figure 2.**
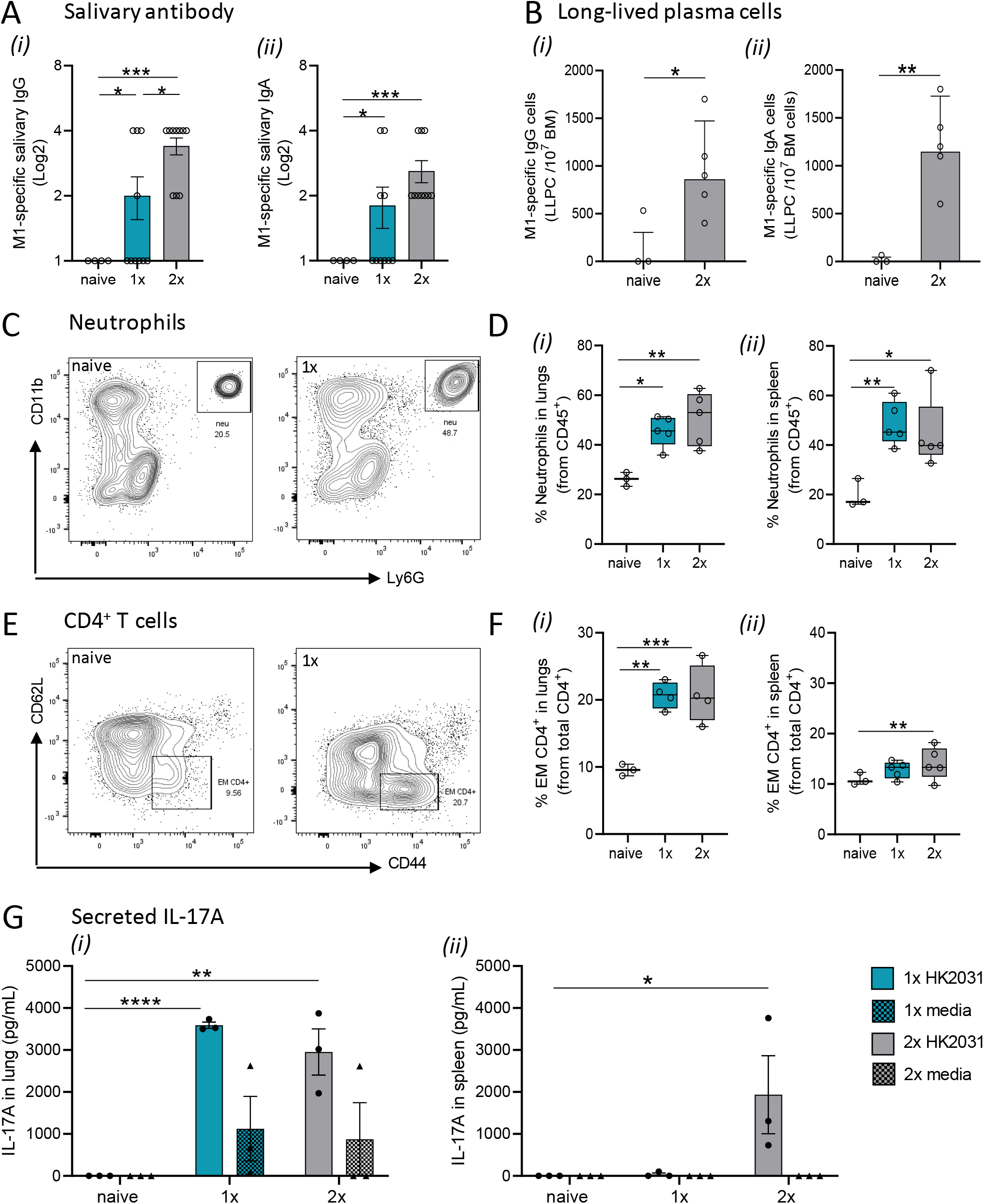
Immune mechanisms involved in mucosal protection. BALB/c mice (n= 5-10 female, 4-6 weeks old) were given a single or 2 IN infections with 2031 (*emm*1), each 3-weeks apart. Three weeks later, all mice received a homologous IN challenge. Mice were sacrificed on day 2 post IN challenge and organs homogenised to produce single cell suspensions for downstream analysis. ***(A) Assessment of salivary IgG and IgA antibody.*** Saliva was collected 7 days following each infection and diluted at 1:2 concentration to determine M1-specific total (i) IgG and (ii) IgA using ELISA. End point titres were defined as the highest dilution of saliva for which the OD was > 3 standard deviations above the mean OD of naive saliva. Significance was determined using One-way ANOVA comparing each group against each other, *p<0.05, *** p<0.001. ***(B) Quantification of long-lived plasma cells in bone-marrow.*** M1-specific (i) IgG and (ii) IgA long-lived plasma cell (LLPC) in the bone marrow were enumerated by ELISpot using 3-5 mice/group and are presented as LLPC per 10^7^ bone marrow (BM) cells. Previously, multiscreen-HA filter plates were coated with 5 μg/mL of M1 extract in carbonate coating buffer overnight. Data shown are geometric mean ± geometric SD minus cell counts from the naive group [to remove background]). Statistical analysis was performed using Mann-Whitney rank analysis, *p<0.05, ** p<0.01. ***(C-D) Neutrophils in lungs and spleen.*** (C) Representative contour plots from flow cytometry analysis of neutrophils (CD45^+^CD11b^+^Ly6G^+^) from the lungs of naïve mice and mice receiving 1 IN infection with 2031. (D) Percentage (%) of neutrophils from CD45^+^ immune cells in the (i) lungs and (ii) spleen is shown. Data are shown as box-and-whisker. ***(E-F) CD4^+^ T cell in lungs and spleen.*** (E) Representative contour plots of CD4^+^ T cell memory populations from the lungs of naïve mice and mice receiving 1 IN infections with 2031. (F) Percentage (%) of effector memory (EM, CD62L^-^CD44^+^) from CD4^+^ T cells (CD3^+^CD4^+^) cells in the (i) lungs and (ii) spleen is shown. Significance was determined using One-way ANOVA comparing each group against each other * p<0.05, ** p<0.01, *** p<0.001. ***(G) IL-17 production by lungs and spleen cells.*** IL-17A responses were assessed in mice that received a single or 2 IN infections. Cell isolates from (i) lungs and (ii) spleens were stimulated *in vitro* with heat killed 2031 (shown as circle) or media as negative control (shown as triangle). At 72 hours post-stimulation, supernatants were isolated, and concentrations of secreted IL-17A was determined using ELISA. Data are presented as pg/mL mean ± SEM. Statistical analysis was performed using Mann-Whitney rank analysis, *p<0.05, ** p<0.01, *** p<0.001. X-axis shows the number of infections prior to challenge. 1x only received infection at week 3 and 2x at both week 0 and 3. Naïve mice are uninfected controls that did not receive infection or challenge.

Next, to explore the role of effector cell-mediated responses in immunity to *S. pyogenes*, we assessed specific cell populations in the lungs and spleen of sequentially infected mice using flow cytometry. Mice that received either 1 or 2 IN infections had a significant increase in Ly6G^+^ neutrophils in both lungs (Fig 2C and 2D i) and spleen (and 2D ii) when compared to naïve mice. Similar observations were made in the lungs and spleen for effector memory (EM) CD4^+^ T cells in mice that received 1 or 2 sequential infections prior to an IN challenge (Fig 2E and 2F).

To assess whether infection-site specific IL-17 secretion was associated with immunity in the URT, we measured IL-17A secreted by lung cells and splenocytes from mice that received 1 or 2 IN infections. For this, immune cells isolated from lung and spleens were stimulated *ex vivo* with heat killed (HK) homologous *S. pyogenes* 2031 prior to detection of IL-17A in culture supernatant by ELISA. Mice that received 1 or 2 infections produced significantly higher (p<0.05) levels of IL-17A in lungs compared to naïve mice (Fig 2G i); however, at least 2 infections were required for induction of IL-17A in spleen (Fig 2G ii).

Taken together these data show that immunity in the URT/LRT following a single *S. pyogenes* IN infection is associated with increase in M1-specific IgG and IgA antibodies in the saliva alongside expansion of neutrophils, EM CD4^+^ T cells and IL-17^+^ cells in the lung.

### IL-17A drives protective immune responses against systemic dissemination following IN infection with *S. pyogenes*

To assess the importance of IL-17A in mucosal immunity against *S. pyogenes* IN infection, we compared IL-17A knockout (IL-17-/-) mice with wild-type (WT) Balb/c mice receiving none or 2 IN infections prior to a final IN challenge (Fig 3A). IL-17-/- mice sequentially infected prior to challenge demonstrated significant increases in bacterial burdens in throat swabs (p<0.01), NALT (p<0.01), lungs (p<0.05), and systemic sites (spleen/lymph nodes, p<0.01) when compared to sequentially infected WT mice (Fig 3B-E). Sequentially infected IL-17-/- mice showed an increasing bacterial burden in the NALT (1.71-fold increase, non-significant), throat swab (3.04-fold increase, non-significant) and systemic sites (4.53-fold increase, non-significant) in relation to naïve (Fig 3B-E). By comparison, sequentially infected WT mice had a significant reduction in bacterial burden in NALT (p<0.05), lungs (non-significant) and in throat swabs (p<0.0001) (Fig 3B-D). Again, as shown previously, the young and immunocompetent mice did not develop systemic infection post challenge (Fig 3E).

**Figure 3.**
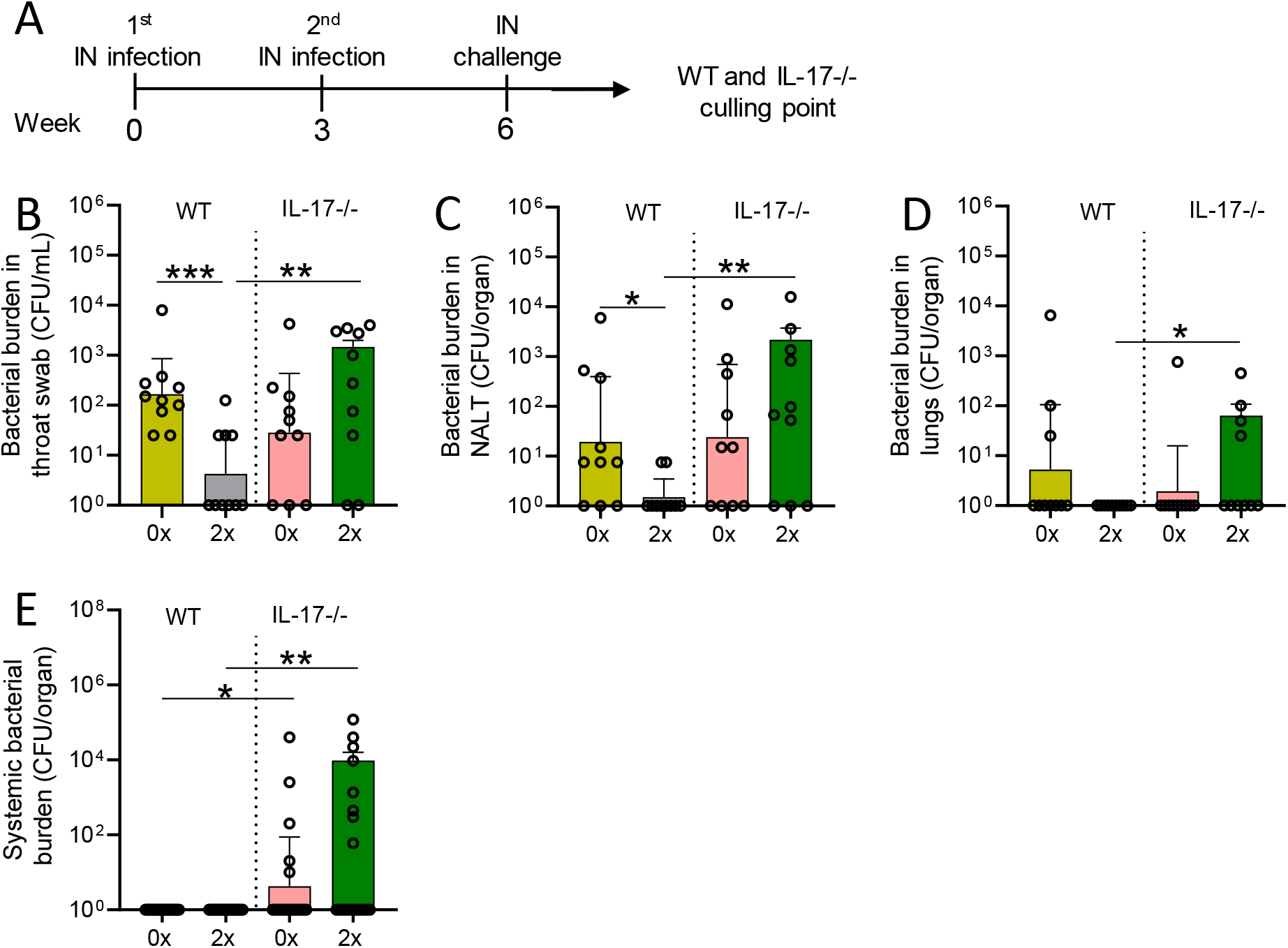
Role of IL-17A in mucosal protection following sequential intranasal infection. (A) IL-17A KO and WT BALB/c mice (n= 10, male and female, 4-6 weeks old) were given 2 IN infections with 2031 (*emm*1), each 3-weeks apart. Three weeks later, all mice received a homologous IN challenge. Mice were sacrificed on day 2 post IN challenge to assess bacterial load in (B) throat swab, (C) NALT, (D) lungs, and (E) systemic sites (spleen and pooled cervical lymph nodes). Data shown are geometric mean ± geometric SD. Significance was determined using Mann-Whitney rank analysis comparing the sequentially infected IL-17-/- mice with their corresponding infected or naïve WT control, * p<0.05, ** p<0.01, *** p<0.001. X-axis shows the number of infections prior to challenge. 2x received infection at both week 0 and 3. 0x received challenge only.

Loss of protection in IL-17-/- mice was associated with a decrease of protective mucosal immune responses against *S. pyogenes.* This was demonstrated by significantly low (p<0.0001) M1-specific IgG antibodies in the saliva of sequentially infected IL-17-/- mice in comparison with WT mice (Fig 4A i). No difference in M1-specific salivary IgA levels was observed between sequentially infected IL-17-/- and WT mice (Fig 4A ii). The number of M1-specific IgG^+^ and IgA^+^ LLPC in BM of IL-17-/- mice that received 2 IN infections, was significantly lower than in WT mice (p<0.05-0.01) (Fig 4B i and ii, respectively). Contrasting with observations in WT mice, M1-specific IgG titres did not change in the sera of IL-17-/- mice after 2 IN infections, (Supp Fig 2A). No changes to IgA responses were observed in the sera of either WT or IL- 17-/- sequentially infected mice (Supp Fig 2B). The role of IL-17 in cell-mediated immunity was assessed in the lungs and spleen of sequentially infected mice. IL-17-/- mice that received 2 IN infections, had a significant decrease in Ly6G^+^ neutrophils in lungs (Fig 4C, D i) paralleled by increase in the spleen (Fig 4D ii), when compared to respective WT control. This shows that mice with intact IL-17A were able to recruit neutrophils from the spleen to the lungs to combat URT/LRT infection.

**Figure 4.**
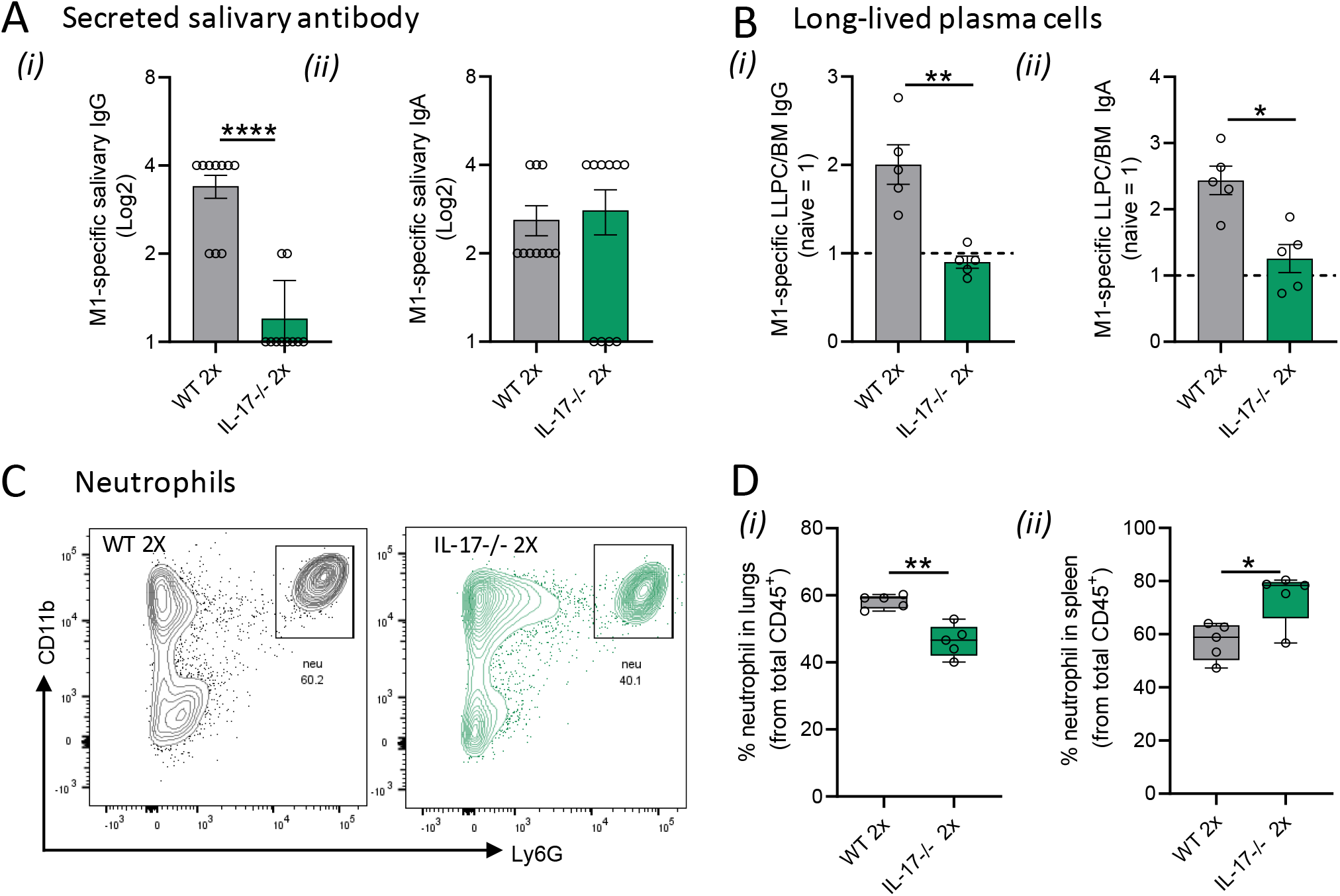
IL-17A-mediated immune mechanisms in mucosal protection. IL-17-/- and WT BALB/c mice (n= 5-10, male and female, 4-6 weeks old) were given two IN infections with 2031 (*emm*1), each 3-weeks apart. Three weeks later, all mice received a homologous IN challenge. ***(A) Secreted salivary antibody in IL-17-/- mice.*** Saliva was collected 7 days following two infections and was diluted at 1:2 concentration to determine M1-specific total (i) IgG and (ii) IgA titers using ELISA. Data shown are antibody titres in saliva from IL-17 2x sequentially infected WT and IL-17-/- mice. ***(B) Quantification of long-lived plasma cells.*** M1 specific (i) IgG and (ii) IgA LLPC in the bone marrow were enumerated by ELISpot using 3-5 mice/group. Data are presented as number of LLPC per 10^7^ bone marrow (BM) cells obtained from 2x sequentially infected WT and IL-17-/- mice ***(C-D) Neutrophils in lungs and spleen.*** Representative contour plots from flow cytometry analysis of neutrophils (CD45^+^CD11b^+^Ly6G^+^) in the lungs of WT mice and IL-17KO mice receiving 2 IN infections with 2031. (D) Data are shown as box-and-whisker plot representing the percentage (%) of neutrophils from CD45^+^ immune cells in the (i) lungs and (ii) spleen from IL-17 2x sequentially infected WT and IL-17-/- mice. Significance was determined using Mann-Whitney rank analysis. * p<0.05, ** p<0.01, comparing naïve IL-17-/- and 2x IN infected IL-17-/- mice.

Taken together, the data show that absence of IL-17 prevents development of natural immunity against *S. pyogenes* in the respiratory tract. This is associated with significantly reduced levels of M1-specific IgG antibodies in the saliva and decreased IgG^+^ and IgA^+^ antibody secreting LLPCs in BM, as well as decreased infiltration of neutrophils and EM CD4^+^ T cells in lungs following infection.

### Repeated IN infections can partially prevent systemic dissemination of *S. pyogenes* after skin challenge

We next modelled a scenario in streptococcal endemic settings to assess if multiple sequential IN infections could provide cross-compartmental protection to the skin and prevent systemic dissemination to lymphoid organs. We previously demonstrated that at least 2 sequential homologous skin infections are required to generate skin immunity against a homologous skin challenge (Pandey et al., 2016a). In the current study, WT mice were given either 2 or 3 homologous IN infections each 3-weeks apart or left naïve (cohort 1). At 3 weeks post last IN infection, all mice received skin challenge with the homologous isolate 2031 (Fig 5A). Six days following skin challenge, mice in each cohort were euthanized to assess bacterial burden in various tissues. We found that none of the mice that received 2 or 3 sequential IN infections were protected from skin challenge (Fig 5B). This was evident from the bacterial burden observed in skin lesions of sequentially infected mice, which were comparable to control mice (Fig 5B). To further investigate if a higher number of prior IN exposures could induce cross-compartmental protection at the skin, a second cohort of mice (cohort 2) received 4 IN infections, each 3-weeks apart, following which, 3 weeks later, they received a skin challenge (Fig 5C). This time, the mice showed a significant reduction (p<0.01) in skin (95%) and systemic bacterial load (95%) when compared to control mice (Fig 5D and 5E, respectively).

**Figure 5.**
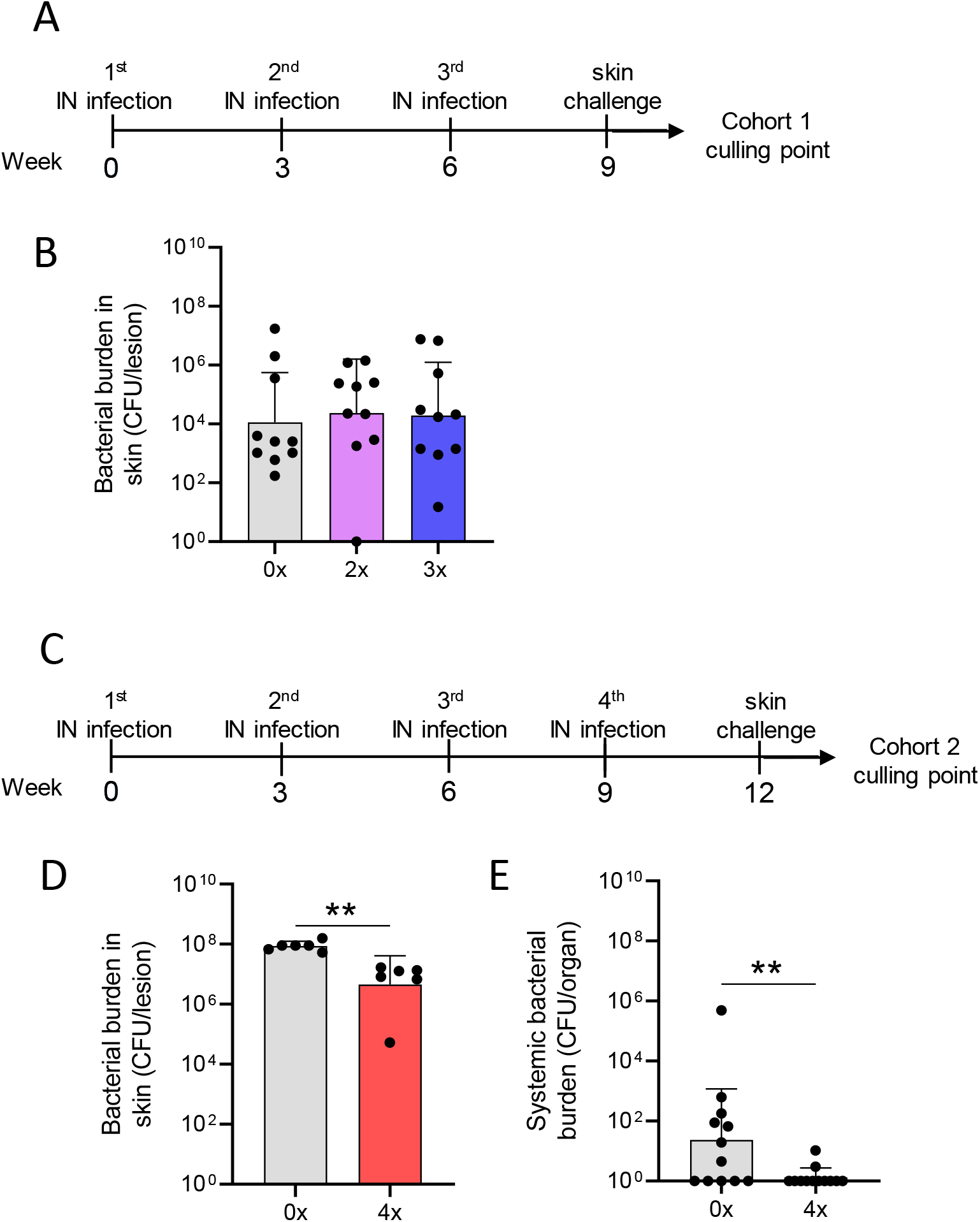
Cross-compartmental protection at the skin following 0-4 prior intranasal homologous infections. (A) Cohort 1 - mice received homologous superficial skin challenge three weeks following 0, 2 or 3 IN infections. BALB/c mice (n= 10, female, 4-6 weeks old) were given IN infections with 2031 (*emm*1), each 3-weeks apart. (B) Mice were sacrificed on day 6 post skin challenge and skin tissues were mechanically homogenised, serially diluted and plated on Columbia blood agar to assess bacterial load (CFU). Skin CFU for the entire skin lesion are presented (C) Cohort 2-mice received 4 IN infections, 3-weeks apart prior to a homologous skin challenge. BALB/c mice (n= 10, female, 4-6 weeks old) were given intranasal (IN) infections with 2031 (*emm*1), each 3-weeks apart. (D) Mice were sacrificed on day 6 post skin challenge to assess bacterial load (CFU) in the skin lesion or systemic sites (represented by combined CFU of spleen and pooled cervical lymph nodes). Skin CFU are calculated for the entire skin lesion, and systemic CFU represents combined CFU of spleen and pooled cervical lymph nodes for each mouse. Data shown are the CFU geometric mean ± geometric SD. Significance was determined using Mann-Whitney rank analysis of CFU in sequentially infected group compared to CFU of the control, ** p<0.01. X-axis shows the number of infections prior to challenge. 2x received infection at both week 0 and 3, 3x at week 0, 3 and 6, and 4x at week 0, 3 and 6 and 9. 0x received challenge only.

Therefore, at least 4 previous IN exposures to *S. pyogenes* at 3 weeks apart are required for development of cross-compartmental homologous protection at the spleen and the skin site.

### Activation of immune responses in the spleen mediate cross-compartmental protection against *S. pyogenes*

To understand whether skin and systemic immunity observed after repeated IN infections were associated with induction of immune responses in the spleen, we analyzed splenic CD4^+^ T cells and neutrophils using immunohistochemistry (IHC) staining. No difference was observed between mice that received 1 or 3 IN infections (Fig 6A and 6B). However, when mice received 4 IN infections, a significant increase (p<0.05) in the number of neutrophils and in the neutrophil elastase H-score (a surrogate marker for neutrophil extracellular trap formation (Yost, Cody et al., 2009)) was observed in comparison to mice that received 1 IN infection (Fig 6A-C). While neutrophils accumulated in the perifollicular spaces, increased numbers of CD4^+^ T cells were observed in the splenic follicles of mice that received 4 IN infections (Fig 6A and D). Interestingly, a significant increase in IL-17 secretion by splenocytes stimulated *ex vivo* with HK *S. pyogenes* 2031 was observed after 3 IN infections; however, it did not correlate with protection from skin challenge (Fig 5B and 6E). CD4^+^ T cell accumulation in the splenic follicles led to the hypothesis that germinal center responses might be playing a role in cross-compartmental protection. To confirm this, we investigated CXCL13 levels in the plasma. CXCL13 is a B cell chemoattractant used as a plasma marker of germinal center activity (Havenar-Daughton, Lindqvist et al., 2016) and also has been shown to play a role in mucosal immunity (Rangel-Moreno, Moyron-Quiroz et al., 2005). We assessed the level of CXCL13 in mouse sera collected at three days and three weeks after each homologous sequential IN infection (Fig 6F). A significant transient increase in serum CXCL13 levels was noted at 3 days post 2^nd^, 3^rd^ and 4^th^ IN infections (4.4 and 5.5-fold increase respectively); however, by 3 weeks post 2^nd^ and 3^rd^ infection the CXCL13 levels had dropped to levels comparable to naïve sera. However, following the 4^th^ IN infection, the significantly increased CXCL13 levels were maintained for at least until 3-weeks post infection (Fig 6F). It is noteworthy that in these mice, a significant increase in CXCL13 levels following 2^nd^ infection also coincided with an increase in serum IgG levels (Supp Fig 1). Although CXCL13 demonstrated a transient, though significant, increase following each IN infection, serum IgG titres remained consistent.

**Figure 6.**
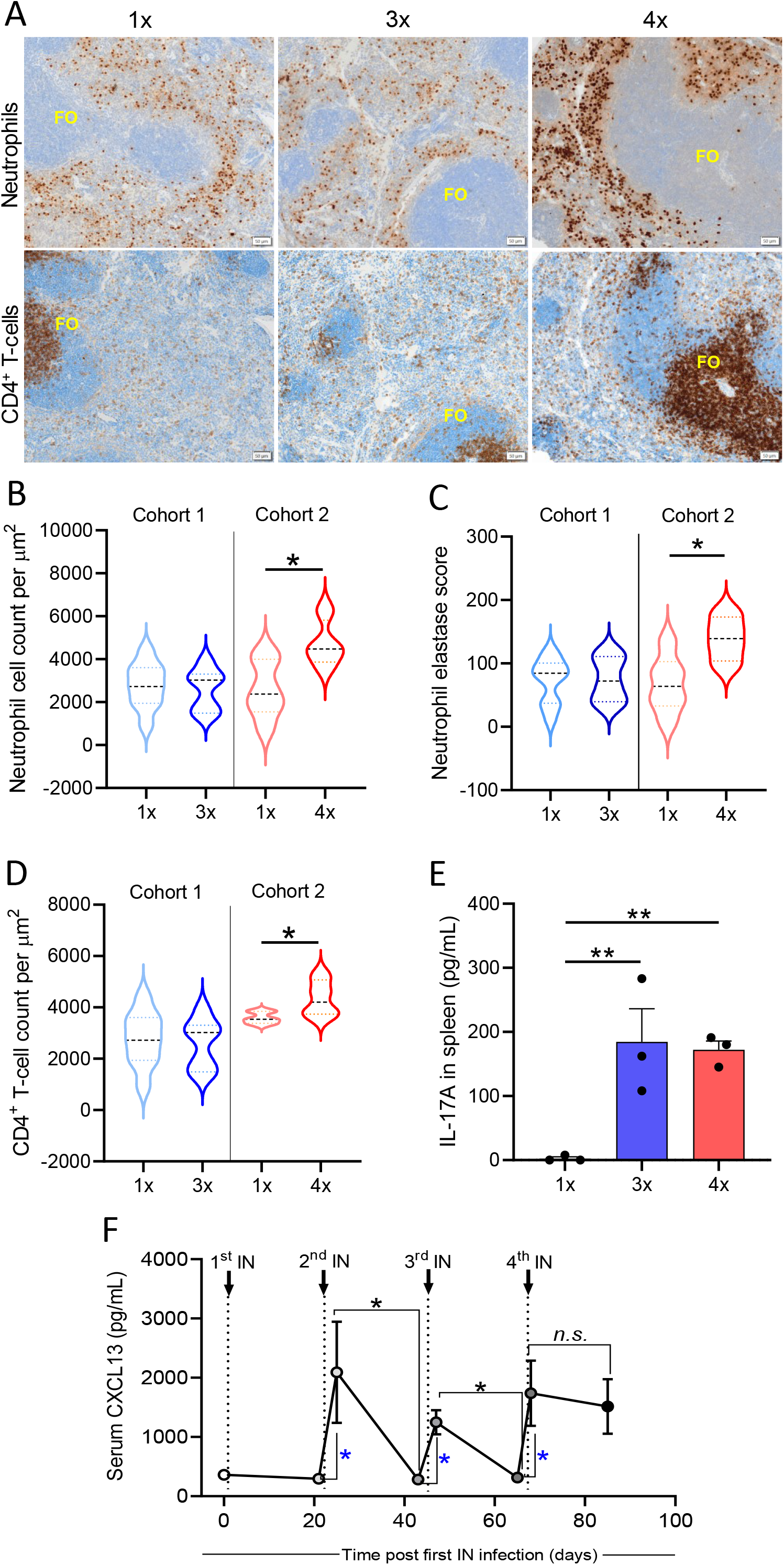
Immune mechanisms associated with cross-compartmental protection. BALB/c mice (n= 5-10, female, 4-6 weeks old) were given repeated IN infections with 2031 (emm1), each 3-weeks apart. Mice were rested for 3 weeks prior to receiving a homologous skin challenge. **(A) Representative immunohistochemistry (IHC) images of neutrophils and CD4^+^ T cells.** Spleen samples from mice with different number of IN infections (1x, 3x, 4x) were formalin-fixed, paraffin-embedded (FFPE) for IHC. Sample were stained with anti-neutrophil elastase and anti-CD4 for the identification of neutrophils and CD4^+^ T cells, respectively, and counterstained with hematoxylin. Splenic follicles (FO) indicated in the figure. Scales bar = 50µm ***(B-D) Enumeration of neutrophils and CD4^+^ T cells.*** (B) Total counts for neutrophils, (C) neutrophil elastase H-score and (D) total counts for CD4^+^ T cells were obtained from entire spleen section. Positive cell detection was used to generate a H-score in QuPath. Cell counts are normalised by total tissue area (um^2^). Data are shown as violin plots depicting distribution of numerical data against number of infections. Cohort 1 received 1 or 3 infections and cohort 2 received 1 or 4 infections. Significance was determined using Mann-Whitney in sequentially infected group compared to 1x infected, * p<0.05. ***(E) IL-17 production by spleen cells*** was assessed using cytokine beads array (CBA). Splenocytes were stimulated *in vitro* during 72h with heat killed 2031 and the supernatant collected for determining the concentrations of IL-17A. Data from duplicates are presented as pg/mL mean + SEM. X-axis shows the number of infections prior to challenge. One Way Anova analysis in sequentially infected group compared to 1x infected, **p<0.01 ***(F) Quantitative assessment of GC response.*** Sera were collected on day 3 and on day 20 post each infection to define concentration of CXCL13. Dotted lines indicate time that 2^nd^, 3^rd^ and 4^th^ IN infections were given. Significance was determined using Mann-Whitney to compare day 3 with day 20 in each infection group, * p<0.05, n.s. = non-significant. Blue * is comparing day 0 (before infection) with day 3 after each sequential infection, * p<0.05. X-axis shows the number of infections prior to challenge. 1x only received infection at week 3, 3x at week 0, 3 and 6, and 4x at week 0, 3 and 6 and 9.

Taken together, repeated URT mucosal infections generated cross-compartmental immunity at the skin and systemic sites that was not associated with changes in IL-17 levels, but rather correlated with neutrophil infiltration and germinal center activity, as evidenced by increased levels of sera CXCL13 and expansion of CD4^+^ T cells in the splenic follicles.

## DISCUSSION

The mechanisms governing natural immunity to *S. pyogenes,* a major human pathogen associated with high morbidity and mortality, are poorly understood due to the pathogen’s diversity and the various infection sites it targets. In this study, we modelled streptococcal exposure in a natural scenario to elucidate the immune mechanisms underlying protection at the respiratory mucosa and cutaneous (skin) sites. Our findings demonstrate a dichotomous role of IL-17 in protection following streptococcal infections. IL-17 played a crucial role in preventing the systemic dissemination of *S. pyogenes* that colonize the respiratory mucosa. Conversely, immunity following skin infection was independent of IL-17 and relied on germinal center formation and neutrophil recruitment.

IL-17 is a proinflammatory cytokine that exerts protective effects against bacterial and fungal infections. It is rapidly released in response to specific triggers at mucosal surface. IL-17 contributes to maintaining epithelial homeostasis, recruiting neutrophils, and stimulating T cell-dependent B cell responses and acts as a crucial link between innate and acquired immune responses (Okada, Puel et al., 2016). Mice deficient in IL-17 receptor have an increased susceptibility to mucoepithelial bacterial infections (Ishigame, Kakuta et al., 2009, Ye, Rodriguez et al., 2001). In this study, we found that deficiency of IL-17 led to *S. pyogenes* escape from the respiratory tract into sterile sites demonstrating its importance in preventing invasive streptococcal disease originating in mucosal sites. This was likely due to inadequate production of M protein-specific IgG antibodies in saliva and impaired recruitment effector immune cells to the respiratory mucosa.

Further, we identified prerequisites for mucosal and systemic protection following repeated IN infections. Previous studies in mice reported the ability to develop mucosal immunity through repeated infections with *S. pyogenes* (Dileepan et al., 2011, Wang, Dileepan et al., 2010). However, it remained unclear what number of infections were required to induce immunity and the duration of protective immune responses. We found that a single mucosal infection occurring 3 weeks prior to a homologous challenge generated *S. pyogenes* type-specific immunity. Given over 250 *emm* types, achieving pan-streptococcal immunity will likely take several years or may never fully occur following natural infection. This explains the high incidence of streptococcal skin and mucosal infections in children under the age of 10, which decreases in early adulthood (Quinn, Vander Zwaag et al., 1985). Furthermore, we have previously demonstrated age-related acquisition of antibodies to the conserved region of *S. pyogenes* M-protein in teenagers and adults residing in endemic areas (Brandt, Hayman et al., 1996), and demonstrated that vaccination of mice with conserved region peptides can induce protection from both mucosal and skin challenge (Ozberk, Reynolds et al., 2021a, Ozberk, Zaman et al., 2023a) (Pandey, Ozberk et al., 2018).

Understanding the mechanisms and longevity of immune responses following infection is of paramount importance for effective public health measures against infectious diseases. Our study investigates immunity at various anatomical sites of *S. pyogenes* infection. Remarkably, we observed that mice developed mucosal immunity after a single IN infection, which persisted for at least 9 weeks. In contrast, our previous findings following S*. pyogenes* skin infection showed that mice required re-infection with the same strain to establish long-lasting immunity (Pandey, Ozberk et al., 2016b). These findings emphasize the existence of distinct immune mechanisms associated with different infection routes. Epidemiological evidence from Australian communities with recurrent pyoderma suggests that repeated skin exposure can confer immunity against throat infections (McDonald, Brown et al., 2007a, McDonald, Towers et al., 2006, Valery, Wenitong et al., 2007). Nonetheless, it is important to note that repeated skin infections have also been implicated in the development of rheumatic fever (Raynes, Frost et al., 2016). Our study aimed to investigate the lack of protection against a single skin infection and assess whether multiple mucosal exposures could provide immunity at the skin and prevent *S. pyogenes* systemic dissemination. We found that mice required a minimum of four homologous mucosal infections to achieve significant protection in the skin and lymphoid organs, underscoring the challenges associated with developing multi-site immunity (Goodfellow, Hibble et al., 2000, McArthur, Medina et al., 2004, Rivera-Hernandez, Pandey et al., 2016).

Only after the fourth IN infection did the levels of CXCL13 become sustained in the plasma. CXCL13 is expressed in lymphoid follicles and acts as a chemotactic signal for CXCR5 expressing B-and T cells (Rangel-Moreno et al., 2005). CXCL13 has been implicated in mucosal immunity by promoting germinal center formation, facilitating Ig isotype switching (Rangel-Moreno et al., 2005), and activating macrophages via IL-17-dependent mechanisms (Gopal, Rangel-Moreno et al., 2013) (Brandt, Teh et al., 2000, Pandey et al., 2016a). Accordingly, we show a modest increase in M-type specific serum IgG after two IN infections, coinciding with the initial increase in CXCL13 systemic levels. Evidence suggest that antibodies produced in response to a primary skin infection can confer type-specific immunity against future streptococcal skin infections (Lancefield, 1959, Pandey et al., 2016a, Wannamaker, Denny et al., 1953). Interestingly, protective immunity in the respiratory mucosa did not rely heavily on germinal center activation or circulating antibodies as mice with low M-specific IgG antibody were still protected after a single infection. Low levels of circulating M-type-specific antibodies during active infection are unsurprising, potentially stemming from IgG cleavage by specific proteases or due to binding to *S. pyogenes* followed by subsequent removal (Frost, Excler et al., 2023). Interestingly, we observed an increase in IgG and IgA secreting cells in the bone marrow following sequential IN infections, aligning with our previous findings that *S. pyogenes* skin infections gradually increased the number of M-type-specific antibody-secreting cells in the bone marrow which led to increased protection (Pandey et al., 2016a).

After the initial priming (first mucosal infection), we observed a significant influx of neutrophils in the lungs which likely led to bacterial killing via antibody-mediated phagocytosis (Andreasen & Carbonetti, 2009). This was paralleled by the increase of lung EM CD4^+^ T cells, which include resident memory T cells that mediate immunity against *S. pyogenes* mucosal infection (Oja, Piet et al., 2018, Ozberk, Zaman et al., 2023b). Lung CD4^+^ T-helper cells have been shown to assist in the development of protective B cell memory responses following respiratory viral infections (Son, Cheon et al., 2021). CD4^+^ T-helper cells producing IL-17 (Th17 cells) effectively support B cell responses and induce pronounced IgG antibody response (Mitsdoerffer, Lee et al., 2010), corroborating our findings of disrupted M-specific IgG antibodies in saliva and BM in IL-17-/-mice. IL-17A is vital for the generation of salivary IgA (Ye, Garvey et al., 2001) and protects against *S. pyogenes* (Carey et al., 2016) and other bacterial infections of the mucosa (Smith, Wasserman et al., 2018) (Chen, Eddens et al., 2016). In the current study, WT mice generated M1-specific antibodies proportionally to the number of mucosal exposures, while IL-17-/- mice had significantly lower IgG antibodies, which correlated with a lack of protection. The protective role of IL-17A through recruitment of neutrophils and tissue-resident memory T cells is evident in our study, as IL-17-/- mice had reduced numbers of lung neutrophils and EM CD4^+^ T cells, as also shown elsewhere (Bagri, Anipindi et al., 2017, Chen et al., 2016, Quinton, Jones et al., 2008). Instead of generating immunity, repeated IN infections to IL-17-/- mice led to *S. pyogenes* accumulation in the respiratory mucosa and invasion of sterile sites. This could be attributed to the fact that in the absence of IL-17, *S. pyogenes* could not be cleared from the respiratory mucosa and accumulated post-each infection. Moreover, IL-17A is vital for maintaining the integrity of epithelial barriers (Song, Zhu et al., 2011).

In summary, our findings highlight that IL-17 orchestrates a multi-faceted mechanism required to generate immunity and prevent systemic dissemination of *S. pyogenes* following mucosal infections. However, it does not prevent *S. pyogenes* infections that start at the skin site. Understanding the molecular and cellular basis of mucosal and cutaneous immune responses will provide important insights into rational design of effective vaccines to prevent superficial and invasive streptococcal disease.

## MATERIAL AND METHODS

### Ethical Statement

Mice were housed at Griffith University’s Animal Facility (Gold Coast, Australia). All experiments and methods for animal procedures were approved by the Griffith University Animal Ethics Committee (Animal Ethics Approval GLY/04/18) in compliance with Australian National Health and Medical Research Council Guidelines. Experimental protocols involving IL-17 knock-out (IL-17-/-) mice were reviewed and approved by Office of the Gene Technology Regulator (OGTR). BALB/c mice (female, 4-6 weeks) were sourced from the Animal Resource Centre, Western Australia. IL-17 -/- mice were acquired from Yoichiro Iwakura (Tokyo University of Science, Japan) (Nakae, Komiyama et al., 2002). Knock-out mice were bred in-house at the Griffith University Animal Facility. Mice were monitored daily for general health. Following challenge, mice were monitored for signs of illness as per a score sheet approved by Griffith University Animal Ethics Committee (Pandey & Good, 2020). Observer blinded to the groups.

### Bacterial strains and culture media

*S. pyogenes* 2031 (*emm*1) was obtained from Menzies school of Health Research (Darwin, NT, AUS). The isolate had previously been serially passaged in mice to ensure virulence and was made resistant to 200 μg streptomycin. The isolate was grown overnight in a liquid Todd-Hewitt broth medium (THB; Oxoid, AUS), supplemented with 1% yeast and 1% neopeptone (THBYN; Difco, AUS). The isolate was 10-fold serially diluted and plated in duplicate on Columbia Blood Agar (CBA; Oxoid, UK), supplemented with 5% defibrinated horse blood (Equicell, AUS) and 200 μg streptomycin (Sigma, China) to determine the number of colony forming units (CFUs).

### Pepsin extraction of the M-protein

The M-protein was extracted using pepsin digestion as described elsewhere (Beachey, Stollerman et al., 1977, Beachey, Campbell et al., 1974). Overnight THBYN cultures were pelleted and resuspended in four-times the weighted volume in PBS pH 5.8 twice. Resuspended pellets were pre-warmed to 37°C for the enzymatic digestion with pepsin A (Merck, AUS) (1 mg pepsin to 10 g bacterial suspension), for 45 minutes with intermittent mixing. The digested suspension was pelleted and the pepM extract buffer exchanged to PBS pH 7 using 10 kDa Amicon centrifugal filter unit (Merck, Ireland). PepM extracts were confirmed with SDS-PAGE and stored in solution at − 20°C.

### Sequential intranasal infection protocol

Mice received sequential intranasal infections 3-weeks apart. Mice were anesthetised via IP injection of ketamine 100 mg/kg and xylazine 20 mg/kg (Zaman, Ozberk et al., 2016). Using a pipette, 5 µL of bacterial inoculum was administered to each nare (1×10^7^ CFU in 10 µL/ mouse), while mouse remained on its back to ensure inhalation. Mice were monitored daily as per above.

### Superficial skin challenge

Mice were challenged using a skin scarification model as previously described (Pandey, Langshaw et al., 2015). Mice were anesthetised with an IP injection of ketamine 100 mg/kg and xylazine 20 mg/kg. The fur from the nape of the neck was removed using clippers and then superficially scarified. A 10 µL inoculum was topically applied, once the inoculum had absorbed into the skin, a temporary cover (Band- Aid™) was applied to the wound and mice were housed individually. Mice were monitored daily for signs of illness as per above.

### Antibody response by indirect Enzyme-linked Immunosorbent Assay (ELISA)

Indirect-ELISA was used to quantify antigen-specific IgG and IgA antibody titres as described elsewhere (Hayman, Brandt et al., 1997). Goat anti-mouse IgG (Bio-Rad, AUS) or or IgA (Invitrogen) horseradish peroxidase (HRP) linked antibodies were used to detect antigen-specific antibodies. Optical density (OD) at 450 nm was measured with a Tecan Infinite m200 Pro plate reader. Titres were defined as the highest dilution of serum for which the OD was > 3 standard deviations (SD) above the mean OD of control samples (naïve sera).

### Sample collection and CFU quantification

Serum samples were collected via puncturing the submandibular vein. Whole blood was allowed to clot and removed prior to centrifuging for sera separation.

Throat swabs were performed using floq swabs (Interpath, USA) moistened in PBS prior to swabbing both sides of the throat. Swabs were squeezed into tubes containing PBS, 10-fold serially diluted and plated onto CBA plates supplemented with 5% horse blood and 200 μg streptomycin. Following overnight incubation at 37°C, CFU were counted to determine bacterial load.

At designated time points, mice were sacrificed via CO_2_ inhalation, whole blood was collected via cardiac puncture into tubes containing Ethylenediaminetetraacetic acid (EDTA; Thermo, AUS). Tissues were collected and mechanically homogenised using a Bullet Blender™ (Next Advance, USA) following the manufacturer’s instructions. Samples were 10-fold serially diluted and plated in replicates onto CBA plates supplemented with 5% horse blood and 200 μg streptomycin. Following overnight incubation at 37°C, CFU were counted to determine bacterial load.

### Cell purification for *in vivo* and *ex vivo* assays

Lymphocyte populations were purified from splenocytes, lung and bone marrow. Lungs were digested in Worthington collagenase III (Scimar, AUS) supplemented with DNase I (Merck, AUS) and all tissues transferred through a 0.70 μM cell strainer (Corning, USA) to obtain single-cell suspensions. RBCs were lysed with ACK lysis buffer and washed with RPMI (Thermo, AUS).

### Antibody-secreting cell response by Enzyme-linked Immunosorbent Spot Assay (ELISpot)

ELISpot was used to quantify the number and location of antibody-secreting cells (ASC) in splenocytes, lung and long-lived plasma cells (LLPCs) in bone marrow. Multiscreen-HA filter plates (Merck, Ireland) were coated with 5 μg/mL of M1 extract in carbonate coating buffer overnight at 4°C. Isolated cells were adjusted to 5×10^6^/mL and directly tested for IgG/IgA-secreting cells using published methods (Pandey, Wykes et al., 2013, Slifka & Ahmed, 1996). Development of spots was performed using AEC substrate kit (BD, AUS) as per manufacturer’s instructions and manually counted to determine ASCs.

### CXCL13 assay

The CXCL13 assay was performed as per manufacturer’s instructions (R&D systems, USA). Test sera was diluted 1:4, and assay standards were prepared in a 1:1 ratio of assay diluent in duplicate and incubated at room temperature for 2 hours on a shaker. Plate was washed 5 times prior to conjugate incubation at room temperature for 2 hours on a shaker. Following washing, substrate solution was added for 30 minutes at room temperature in the dark, prior to addition of stop solution. OD at 450 nm was measured with a correction of 540 nm, with a Tecan Infinite m200 Pro plate reader.

### IL-17A ELISA

The IL-17A ELISA was performed as per manufacturer’s instructions (Mouse IL-17A ELISA MAX; Biolegend, USA). Nunc MaxiSorp plates (Thermo, AUS) were coated overnight with the IL-17A capture antibody diluted in a carbonate coating buffer. Plates were blocked with 1% BSA/PBS for 1 hour prior to washing. Splenocyte supernatants (diluted 1:4, in duplicate) and standards were incubated at room temperature for 2 hours. Detection antibody was incubated for 1 hour prior to avidin-HRP incubation for a further 30 minutes. TMB substrate solution for was incubated for 20 minutes, and then the addition of an acid stop solution. Absorbance (OD) was read at 450 nm with a correction of 570 nm. IL-17A concentration was determined by plotting unknown samples on a standard curve to determine IL-17A pg/mL.

### Cytokine production by Cytometric bead Analysis (CBA)

Single cell suspension of splenocytes were prepared as above, counted using a haemocytometer in trypan blue (Sigma, AUS) and adjusted to 4×10^6^ cells per mL. Splenocytes were stimulated with concanavalinA (ConA), lipopolysaccharide (LPS), pepsin M1 extract, heat-killed 2031 or media as a negative control. After 72-hour stimulation at 37°C with 5% CO2, cells were centrifuged at 300 g for 10 minutes and supernatant collected. Supernatants were stored at −80°C. Cytokines in supernatant were subsequentially measured using a mouse Th1/Th2/Th17 CBA kit (BD, AUS) as per manufacturer’s instructions. Samples were acquired using the BD LSR Fortessa cytometer, and data analyzed using FCAP Array v3.1 (BD).

### Cell population analysis by Flow Cytometry

Cell populations were determined by flow cytometry. Single cell suspensions were prepared as described above and were pre-incubated with Fc block (CD16/32) for 15 minutes on ice. Cells were surface stained with a master mix containing a dead cell exclusion dye (NIR), CD45-BUV395, CD11b-BUV737, CD3-PE-CF594, and Ly6G-BV510. Cells were stained on ice in the dark, for 40 minutes. Following incubation, cells were washed in FACS buffer (2.5% fetal calf serum, 5mM EDTA in PBS) and fixed in 2% paraformaldehyde for 15 minutes. Samples were washed in PBS and acquired on a BD LSR Fortessa flow cytometer, and data analyzed using FlowJo V.10.7 (BD).

### Immunohistochemistry (IHC)

Formalin-fixed paraffin-embedded (FFPE) spleens were sectioned at 3 μm on SuperFrost+ slides. IHC staining of spleen sections for CD4 (1:200, D7D2Z; Cell Signaling Technology) and neutrophil elastase (1:200, E8U3X; Cell Signaling Technology) was performed on the Leica BOND™ RX auto-stainer (Leica, Nussloch, Germany) using the BOND Polymer Refine Detection (Leica) kit and developed with 3,3’-diaminobenzidine as the chromogen. Stained slides were mounted in Dako Mounting Medium (Dako,) and coverslipped using a Dako coverslipper.

### IHC acquisition and analysis

Images were acquired using an Olympus VS200 digital slide scanner (EVIDENT Life Science, USA) under the brightfield emission. Each tissue section was acquired using a 20× objective (UPLXAPO 20X; NA 0.8). Individual images were obtained from regoins of interest (ROI) drawn using the VS200 software based on automatic outline thresholding. Cell counts were then performed using QuPath (v 4.0.1), and neutrophil elastase H-score was assigned in the range of 0-300. Cell counts are given as average of cells per µm2Data was exported to Microsoft Excel for tabulation and plotted in GraphPad Prism.

### Statistical analysis

Data were analyzed using Graph Pad PRISM version 10.7 for Windows. All data, except where noted, are represented as the geometric mean <u>±</u> standard error of the mean (SEM). Statistical differences between two groups were determined using non-parametric Mann-Whitney *t*-test. When comparing more than two groups, analyses were performed using one-way ANOVA with *p*<0.05 considered to be statistically significant.

## Data availability

The authors declare that all data supporting the findings of this study are available within the paper and its supplementary information.

## ACKNOWLEDGEMENT

This work was supported by the National Health and Medical Research Council, Australia (NHMRC) Project Grant (APP1160379), Program Grant (APP1083548), and Investigator Fellowship (APP1174091) to MFG. J-L.M is supported by Australian Postgraduate Award, Australian Government and Griffith University.

